# Decrypting the phylogenetics history of EGF-CFC proteins Cripto and Cryptic

**DOI:** 10.1101/2024.08.30.610562

**Authors:** Natalia A. Shylo, Paul A. Trainor

## Abstract

EGF-CFC proteins are obligate coreceptors for Nodal signaling and are thus required for gastrulation and left-right patterning. Species with multiple family members show evidence of specialization. For example, mouse *Cripto* is required for gastrulation, whereas *Cryptic* is involved in left-right patterning. However, the members of the family across model organisms have little sequence conservation beyond the EGF-CFC domain, posing challenges for determining their evolutionary history and functional conservation. In this study we outline the evolutionary history of the EGF-CFC family of proteins. We traced the *EGF-CFC* gene family from a single gene in the deuterostome ancestor through its expansion and functional specialization in tetrapods, and subsequent gene loss and translocation in eutherian mammals. Mouse *Cripto* and *Cryptic*, zebrafish *Tdgf1,* and all three *Xenopus EGF-CFC* genes (*Tdgf1*, *Tdgf1.2* and *Cripto.3*) and are all descendants of the ancestral *Tdgf1* gene. We propose that subsequent to the family expansion in tetrapods, *Tdgf1B* (*Xenopus Tdgf1.2*) acquired specialization in the left-right patterning cascade, and after its translocation in eutherians to a different chromosomal location, *Cfc1/Cryptic* has maintained that specialization.

## INTRODUCTION

Nodal signaling pathway is central to body patterning due to its involvement in gastrulation, anterior-posterior, dorsal-ventral and left-right patterning processes^1^. EGF-CFC proteins are membrane-anchored and serve as obligate coreceptors for Nodal to assemble the receptor complex, thus making them indispensable for most aspects of Nodal signaling – mesoderm and endoderm formation, as well as anterior-posterior and left-right patterning^2,3^. The family of EGF-CFC proteins got its name from a conserved domain, composed of an EGF-like motif, and a novel <u>CFC</u> sequence, first identified in mouse <u>C</u>ripto, frog <u>F</u>RL-1, and mouse <u>C</u>ryptic/Cfc1^4^. Zebrafish Tdgf1 is another well-known member of the family^5^. *one-eyed-pinhead (oep*) mutation in zebrafish *Tdgf1* is frequently used to knock out all Nodal signaling^6^. Despite high conservation of the EGF-CFC domain, the protein sequence is poorly conserved amongst members of the family. This led to hypotheses that there was no orthologous relationship between family members from different species^4,7,8^.

The mouse genome possesses two *EGF-CFC* genes – *Cripto* and *Cryptic*, which have specialized functions. *Cripto* regulates mesoderm formation and anterior-posterior patterning, whereas *Cryptic* influences left-right patterning^9^. In contrast, single zebrafish *Tdgf1* fulfils both functions of mesoderm formation and left-right patterning^10^. In humans, *Cryptic*/*CFC1* underwent an additional duplication, resulting in *CFC1B* and *CFC1* genes (Figure 3B) and mutations in both genes have been reported in patients with congenital heart disease and heterotaxy, suggestive of roles in left-right patterning^11,12^.

The evolutionary history of the *EGF-CFC* gene family is heretofore unknown, and orthologous relationships between the members of the family remain contentious. It has been suggested that *EGF-CFC* genes underwent a duplication event after the speciation of vertebrates from chordates, and that the gene corresponding to the *Cripto* lineage was subsequently lost in non-mammalian species, while being retained and conserved in eutherian mammals^2,8^.

With the recent exponential growth of sequenced and annotated genomes, we revisited the evolutionary history of the EGF-CFC family and discovered that the *EGF-CFC* genes indeed underwent a duplication event as part of a larger gene cassette, after the speciation of vertebrates. However, our analysis shows that one of the two orthologous genes was lost shortly after the duplication event. A more functionally significant event was gene expansion in a tetrapod ancestor that resulted in three tandem copies of *Tdgf1* gene (named *Tdgf1A-C* here). Based on protein conservation we have determined that *Tdgf1A* was lost in the eutherian ancestor, and that *Tdgf1B* was translocated to a novel chromosomal position and is now known as *Cryptic/Cfc1*. Considering that the *Xenopus Tdgf1B* ortholog *Tdgf1.2* is involved in left-right patterning similar to *Cryptic/Cfc1*, we propose that this specialization occurred in a tetrapod ancestor and has been conserved ever since.

## RESULTS

### *EGF-CFC* genes in deuterostome genomes

The evolutionary history of EGF-CFC protein family has been previously examined in a limited capacity, mostly restricted to the sequence comparison of the family members to determine their evolutionary relationship^2,8,13^. With vastly more genomes now available, we carried out a thorough survey of the *EGF-CFC* genes, present in deuterostome genomes, and corroborated their orthologous relationships through analysis of regions of chromosomal homology and sequence conservation.

This survey of the *EGF-CFC* genes in deuterostomes revealed that all deuterostomes, except for tunicates (ex. *Cional intestinalis*) have at least one copy of an *EGF-CFC* gene, with the ancestral gene most frequently named *Tdgf1 (teratocarcinoma-derived growth factor 1)* (Figure 1). Furthermore, most eutherians have an additional *Cryptic/Cfc1* gene, except for the cow genome in which it has been lost, and in humans, which experience a duplication in that region, resulting in two genes – *CFC1B* and *CFC1*. We noted additional gene expansions in amphibians and reptiles in the *Cripto/Tdgf1* region, with many lineages having 2-3 genes in the same chromosomal region (Figure 1).

**Figure 1.**
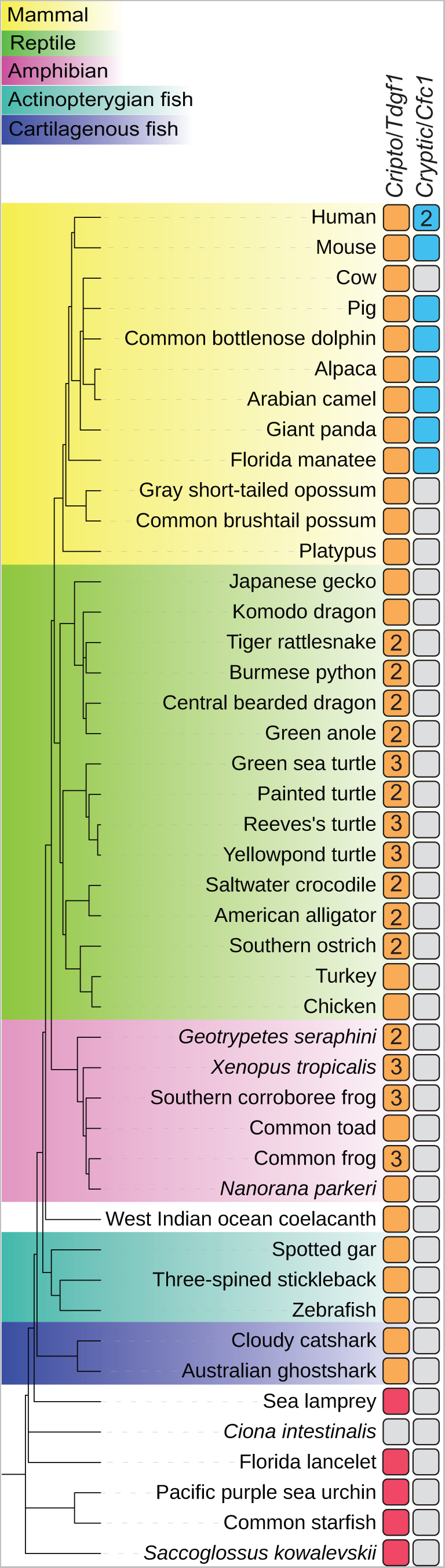
| Phylogenetic distribution of EGF-CFC proteins. The phylogenetic relationship of deuterostome clades evaluated in this study, showing presence or absence of *Crypto/Tdgf1* and *Cryptic/Cfc1* homologs in their genomes. The numbers in the squares indicate additional expansion in that region and the number of genes. Orange indicates the *Cripto/Tdgf1* gene is present in jawed vertebrates, whereas red is indicative of the ancestral gene. *Crypto/Cfc1* is demarcated with a blue colored square.

### Ancestral *Tdgf1* gene

The key to determining homologous relationships between genes is examining them in the context of gene linkage groups across the phylogenetic tree. Although the *EGF-CFC* family originated in a urbilaterian ancestor^2^, in this study we limited our analysis to deuterostomes. Using previously published protein sequences^2^, we began by identifying the *Tdgf1* genes in echinoderms, hemichordates and chordates like the Florida lancelet (Supplementary Table 1). Although we noted several rearrangements and recombinations, we also identified clear linkage groups (Figure 2). Most importantly, *Tdgf1* is linked to *Rtca* and *Dbt1* in echinoderms, hemichordates and lancelets (Figure 2). Although we were able to locate *Rtca, Dbt1* and other genes from the linkage group in *Ciona intestinalis*, we were unable to identify *Tdgf1* in the genome, consistent with prior observations^2^. The *Tdgf1* gene in the craniate sea lamprey was located at a novel chromosomal position, however, our analysis was complicated by the limitations of partial genome assembly^14^.

**Figure 2.**
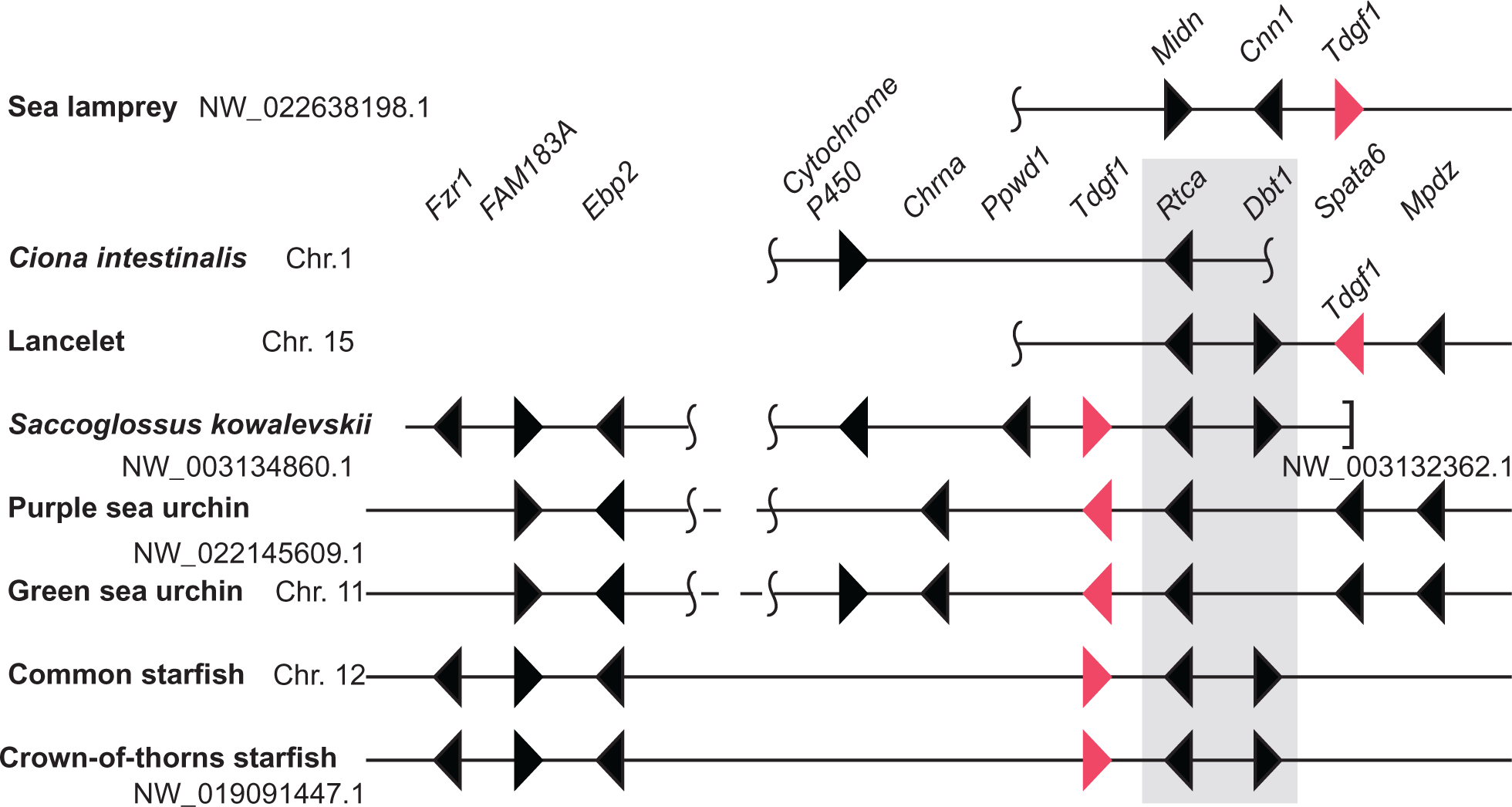
| Structure of the chromosomal regions containing the ancestral *EGF-CFC* gene *Tdgf1* in deuterostomes. The ancestral *Tdgf1* gene is demarcated by a red triangle. Expanded chromosomal regions and select landmark genes for lancelet, green sea urchin and common starfish are in figure S1B. *Dbt* and *Rtca* are linked to *Tdgf1* on the ancestral deuterostome chromosome and are highlighted in gray. Vertical tildes designate the location of an ancestral chromosomal recombination events. A square bracket represents the edge of a genomic scaffold. Dashed lines between two sections indicate that the genes are on distant parts of the same chromosome/scaffold. Chromosomes and scaffold numbers are as indicated. Only conserved landmark genes are pictured. Additional species-specific genes are present throughout, and not depicted. Pictured are key model organisms, and additional species, which aid in revealing conserved regions of homology.

### *EGF-CFC* genes in jawed vertebrates

We then tracked the genomic history of *Tdgf1* further into the jawed vertebrate lineage. Although we detected linkage between *Lrrc2, Cripto/Tdgf1* and *Fam240* in most jawed vertebrates we examined, which clearly established orthologous relationship between these genes (Figure 3A), this linkage group had no apparent similarity with the ancestral *Tdgf1* region (Figure 2), and the linkage between *Tdgf1, Dbt1* and *Rtca* was apparently lost.

**Figure 3.**
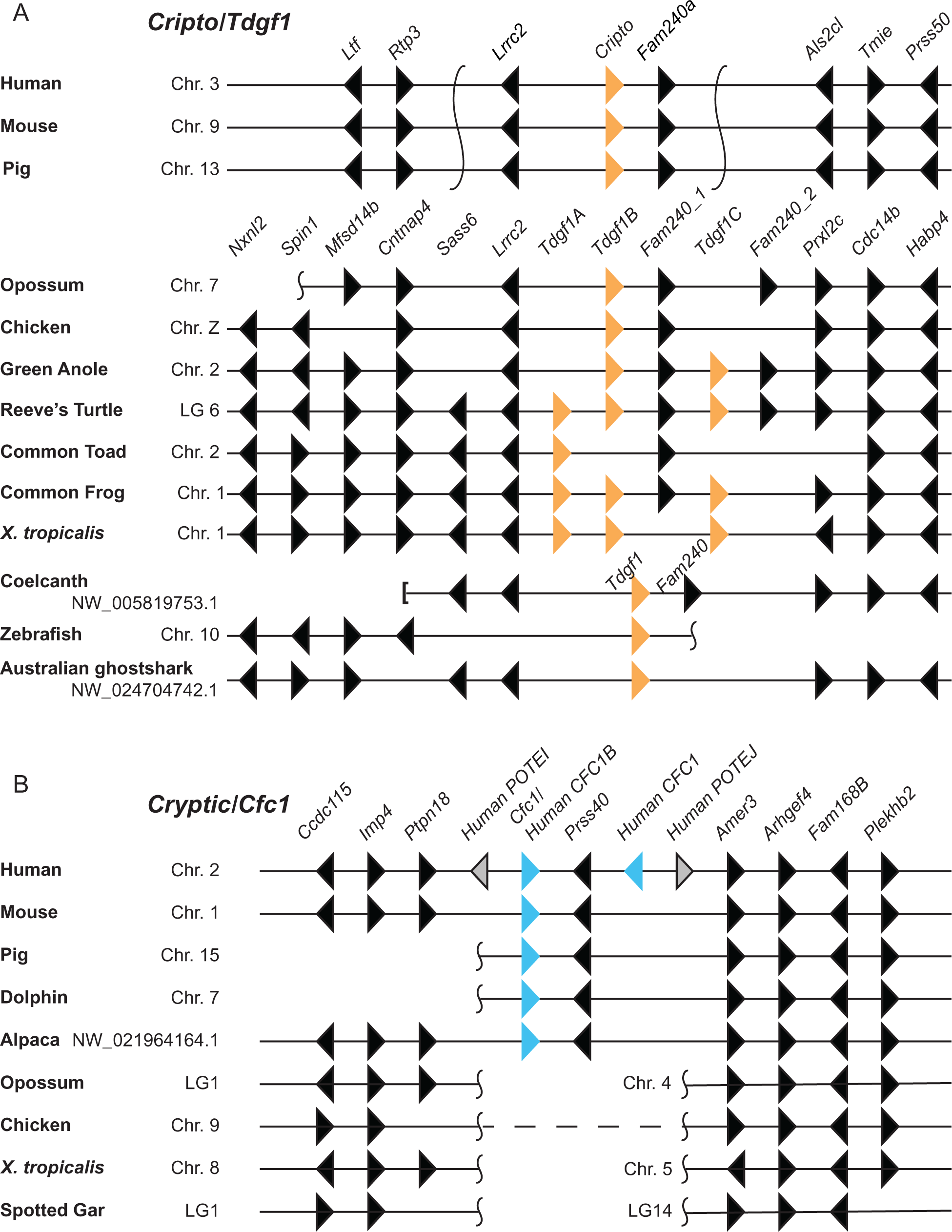
| Structure of the chromosomal regions containing *EGF-CFC* gene family members in jawed vertebrates. A. The *Cripto/Tdgf1* present in jawed vertebrates is demarcated by orange triangles. This gene underwent an expansion in tetrapods. We termed the resulting three genes *Tdgf1A*, *Tdgf1B* and *Tgf1C* in order from *Lrrc2*. Duplicated *Fam240* genes were labeled *Fam240_1* and *Fam240_2* in order from *Lrrc2*. **B.** Structure of the chromosomal regions containing *Cryptic/Cfc1* genes, demarcated by blue triangles. Only conserved landmark genes are pictured. Additional species-specific genes are present throughout, and not depicted. Pictured are key model organisms, and additional species, which aid in revealing conserved regions of homology. Vertical tildes designate the location of an ancestral chromosomal recombination events. A square bracket represents the edge of a genomic scaffold. Dashed lines between two sections indicate that the genes are on separate distant parts of the same chromosome. Chromosomes and scaffold numbers are as indicated.

Interestingly, in jawed vertebrates *Dbt1* and *Rtca* remain linked, but lack an *EGF-CFC* gene nearby (Supplementary Figure S1A). We noted the similarity between the extended genomic region containing *Dbt1* and *Rtca* and the one containing *Cripto/Tdgf1* in jawed vertebrates, most notably the presence of *Lrrc* family genes in proximity to *Tdgf1* and *Dbt1* (*Lrrc2* and *Lrrc39*, respectively), indicating that these regions likely arose as the result of a duplication, with subsequent gene loss (Figure 3A, Supplementary Figure S1A). This likely happened in a vertebrate ancestor, since only a single region of homology is present in lancelets (Supplementary Figure S1B), and only single copies of genes from the linkage groups could be identified in *Ciona intestinalis* (not shown).

However, both regions (Figure 3A and Supplementary Figure S1A) are still markedly different from the relatively small region of homology that contains the ancestral *Tdgf1* that we highlighted (Figure 2). An extended analysis of landmark genes in the lancelet, green sea urchin and common starfish, spanning larger chromosomal regions (Supplementary Figure S1B) reveal that the genes present in the two duplicated regions in jawed vertebrates (Figure 3A, Supplementary Figure 1A) are all present on a single chromosome in those deuterostomes, with a clearly identifiable linkage block intercalated with each other (Supplementary Figure S1). Therefore, mouse *Cripto*, zebrafish *Tfgf1,* and all three *Xenopus EGF-CFC* genes (*Tdgf1*, *Tdgf1.2* and *Cripto.3*) and are all descendants of the ancestral *Tdgf1* gene.

This analysis didn’t, however, reveal the origin of the *Cryptic/Cfc1* locus. It emerged in eutherian mammals, and gene linkage analysis of the region did not reveal any homology to *Cripto/Tdgf1* and the ancestral *Tdgf1* regions.

### *Tdgf1* gene expansion

We next examined more closely the *Tdgf1* genomic region in amphibians and reptiles. *Xenopus Frl-1* (*Cripto.3*) is one of the founding members of the *EGF-CFC* family and one of three *Tdgf1* genes in *Xenopus*^7^. *Xenopus* is the only widely used model organism that has three *Tdgf1* genes (Figure 1). We will refer to them as *Tdgf1A*, *Tdgf1B* and *Tgf1C*, corresponding to *Xenopus Tdgf1*, *Tdgf1.2* and *Cripto.3*, accordingly (Figure 3A). Humans, mice, chickens and zebrafish all contain a single copy of the *Cripto/Tdgf1* gene in that genomic locus (Figure 1), so it was reasonable to assume that this gene expansion was unique to *Xenopus*, similar to the expansion of *Nodal* genes to over 7 in some frog species^15,16^.

However, our analysis of amphibian species revealed other lineages with 2 or 3 *Tdgf1* genes, like the caecilian *Geotrypetes seraphini* (2 genes), and common frog (3 genes) (Figure 1, Supplementary Table 1), with *Fam240* located between *Tdgf1B* and *Tdgf1C*. Notably, we identified many reptilian species also containing 2-3 *Tdgf1* genes, e.g. tiger rattlesnake (2 genes), green sea turtle (3 genes), and painted turtle (2 genes) (Figure 1, Supplementary Table 1), some of which also contain one or two *Fam240* genes interspersed with *Tdgf1* genes (we will refer to these as *Fam240_1* and *Fam240_2*) (Supplementary Figure S2A). Based on the similarity of their positioning in the genomic locus, we wondered if the three *Tdgf1* genes and two *Fam240* genes in amphibians and reptiles were orthologous. Thus, we compared the sequences of a large number EGF-CFC proteins, uncovering an orthologous relationship between the three *Tdgf1* genes in amphibians and reptiles (Figure 4). Likewise, *Fam240* genes show clear orthology in the species examined (Supplementary Figure S2B).

**Figure 4.**
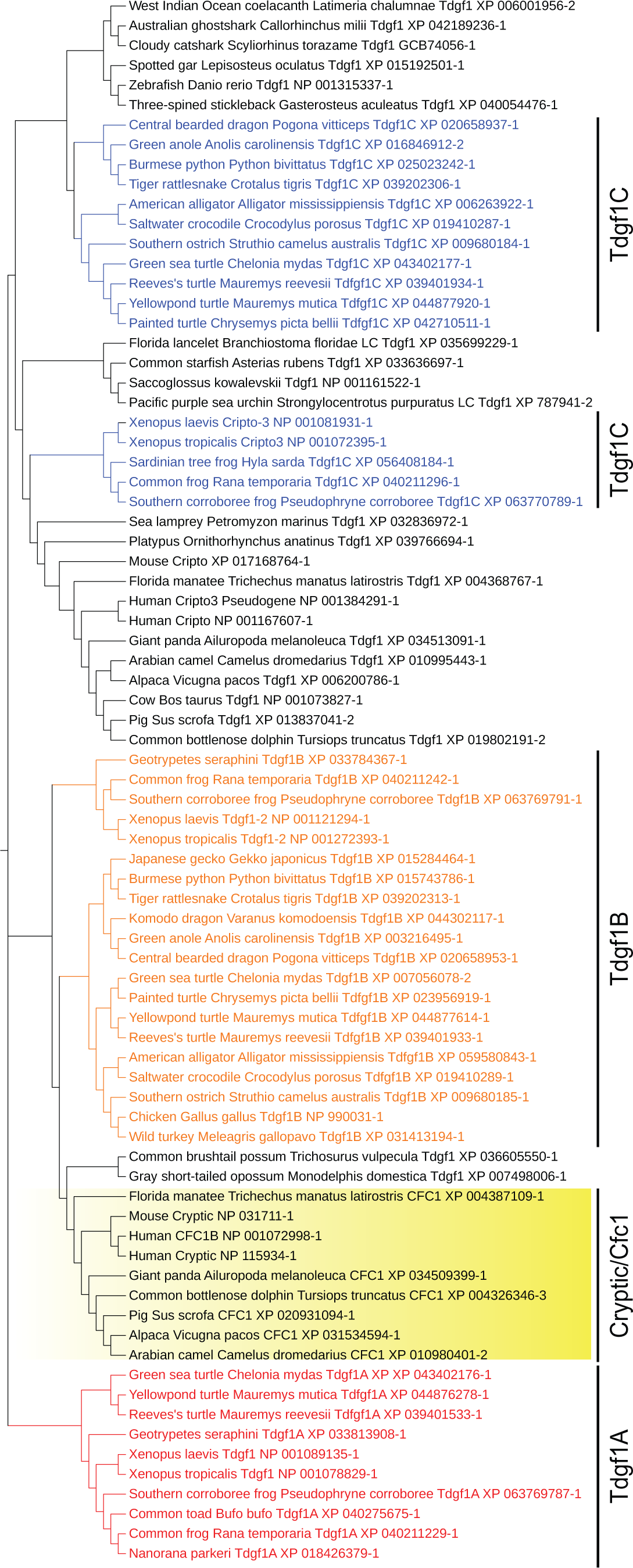
| Phylogenetic relationship of EGF-CFC proteins. The phylogenetic relationship of EGF-CFC protein sequences in deuterostomes. Cryptic/Cfc1 in eutherian mammals is highlighted in yellow text. Tdgf1A proteins are in red, Tgf1B sequences are in orange, and Tdgf1C are in blue.

The protein clustering revealed patterns of gene inheritance (Figure 4). We found that Tdgf1A is restricted to amphibians and reptiles, whereas Tdgf1C is closely homologous with the eutherian Cripto/Tdgf1 proteins. Notably, Tdgf1B is most closely related to Cryptic/Cfc1 protein, finally revealing its evolutionary origin (Figure 4). These data suggest that *Tdgf1B* underwent translocation to a new locus in the genome and is now known as *Cryptic/Cfc1* (Figure 5). Interestingly, platypus Tdgf1 is most closely related to Tdgf1C, whereas marsupial Tdgf1 protein is more closely related to Tdgf1B. Platypus has retained *Tdgf1C*, whereas marsupials have retained *Tdgf1B*, and the eutherian mammals have retained *Tdgf1C* in the ancestral locus, with a *Tdgf1B* translocation to a novel genomic locus, where it became known as *Cryptic/Cfc1*. With additional evidence from two *Fam240* genes in the marsupial genomes, and their position in relation to *Tdgf1* gene, we propose that the last common eutherian ancestor had two *Tdgf1* genes (*Tdgf1B* and *Tdgf1C*) and two *Fam240* genes.

**Figure 5.**
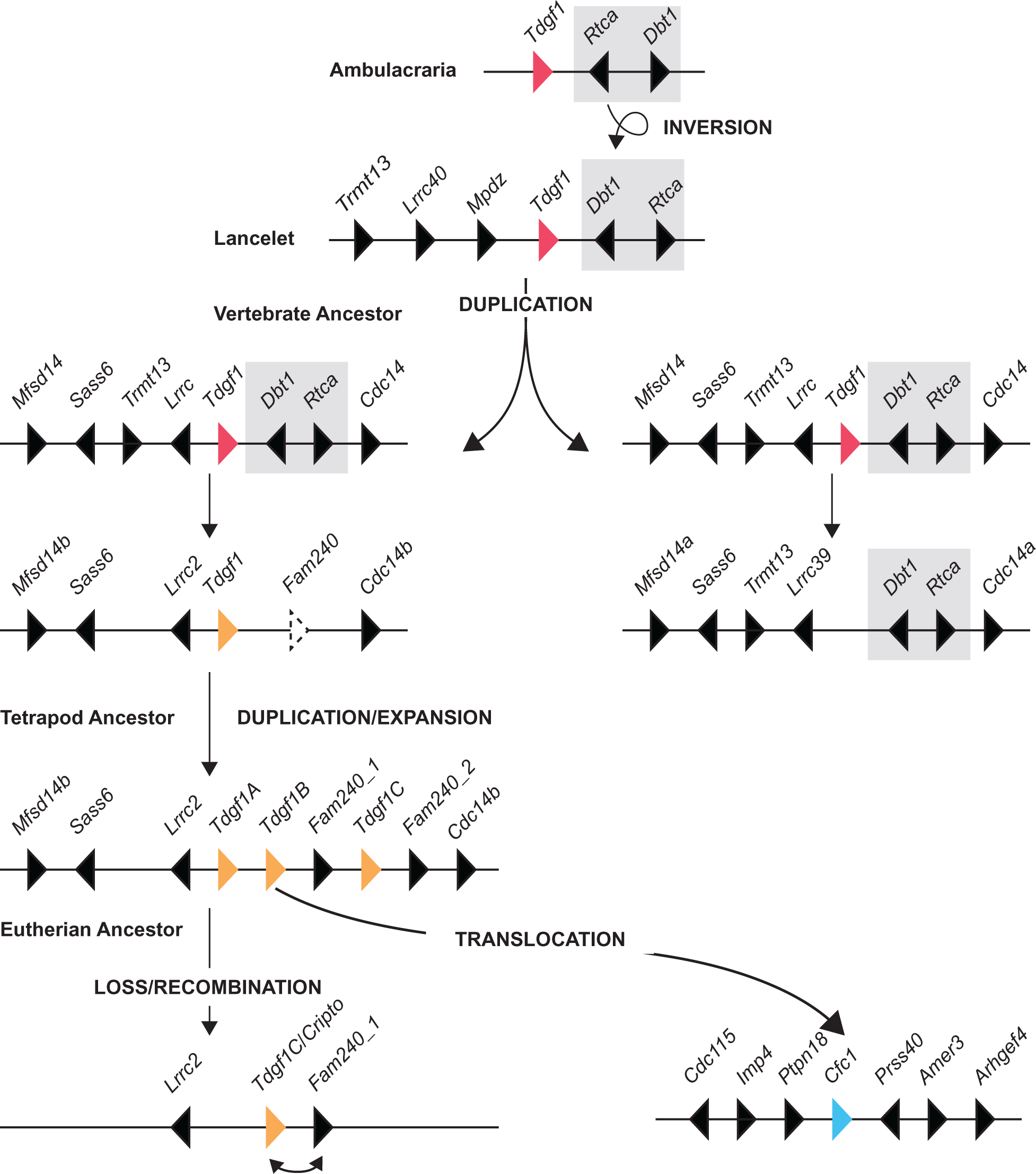
| Model of the *EGF-CFC* genomic regions evolution. *Dbt* and *Rtca* genomic cassette, adjacent to the ancestral *Tdgf1* gene, underwent an inversion in a chordate ancestor. In a vertebrate ancestor the entire genomic region underwent duplication, followed by gene loss in both regions. We also note acquisition of *Fam240* gene, proximal to *Tdgf1*, somewhere in the fish lineage. In a tetrapod ancestor, the *Tfgf1* and *Fam240* genes underwent an expansion event, resulting in three copies of *Tdgf1* (named here A-C, counting from *Lrrc2* gene) and two copies of *Fam240* (named here 1 and 2, counting from the *Lrrc2* gene). Subsequently in the eutherian ancestor the ancestral region including *Tdgf1A* was lost, and *Tdgf1B* underwent translocation to a new genomic site, becoming known as *Cryptic/Cfc1*. The ancestral *Tdgf1* gene is demarcated by a red triangle, whereas *Cripto/Tdgf1* is demarcated in orange and Cryptic/*Cfc1* is light blue.

## DISCUSSION

Here we report on the evolutionary history of the EGF-CFC family of proteins, which act as obligatory co-receptors for Nodal signaling. A newcomer to the field of Nodal-related research may find it difficult to unravel the relationships between the genes in the *EGF-CFC* family, as it may be not immediately clear what the difference is between Cripto and Cryptic, and how they relate to Cfc1 and Tdgf1 proteins. In addition, which model species have a single *EGF-CFC* gene, which ones have more, and what may be the difference in their functions, has yet to be succinctly described and summarized. Furthermore, in addition to a collection of historic names and several acceptable aliases, gene names have not been assigned altogether to the identified transcripts in nearly half of the genomes examined in this study, making any work with this gene family particularly challenging.

Previous studies revealed that the *EGF-CFC* gene first appeared in the ancestor of all bilaterians – an urbilaterian and can be found in deuterostomes and several lophotrochozoans^2,17^. Interestingly, no *EGF-CFC* genes could be identified in ecdysozoans, which have also lost the *Nodal* gene, thus suggesting they co-evolved^2^. Several studies in deuterostomes such as tunicates, also failed to identify an *EGF-CFC* transcript, and our analysis in *Ciona intestinalis* of the genomic regions, linked to *Tdgf1* in other animals, has confirmed its absence in the genome (Figure 1, Supplementary Table 1). *Ciona intestinalis* does, however, have *Nodal*, and given the requirement for EGF-CFC proteins for proper Nodal signaling, the mechanism for Nodal signal transduction in tunicates begs to be investigated.

Our analysis did however reveal a single *EGF-CFC* gene (*Tdgf1*) in echinoderms and hemichordates, as well as the Florida lancelet, linked to *Rtca* and *Dbt1* genes (Figure 2, 5). Subsequently, a large region containing *Tdgf1* underwent duplication (Figure 5). Shortly thereafter genes became lost from the two loci, such that *Cripto/Tdgf1* gene was lost from the proximity of *Rtca* and *Dbt1* in one duplicated region, and *Rtca* and *Dbt1* were lost from the other region (Figure 5). We have not yet identified a genome where *Tdgf1* is present in both duplicated regions, pointing to rapid gene loss from the two regions.

We propose that the duplication occurred in an early vertebrate ancestor, since we don’t find evidence for it in tunicates, but we did detect some duplicated cassettes in the sea lamprey. However, the current state of genome assemblies and annotation for *Ciona intestinalis* and lampreys make a more precise analysis challenging.

Arguably, the most significant event in the history of *EGF-CFC* genes occurred in a tetrapod ancestor, when the *Tdgf1* gene underwent expansion, resulting in three copies of *Tdgf1* in tandem (Figure 5, Supplementary Figure S2A). Subsequent loss of *Tdgf1* paralogs in different tetrapod lineages obscured the orthologous relationship between genes, making it difficult to unravel these relationships until now. In eutherian mammals *Tdgf1A* was entirely lost from the genome, and *Tdgf1B* underwent a translocation, coupled with chromosomal rearrangements. Genomic regions previously located on separate chromosomes came together, with *Tdgf1B* landing between them, becoming known as *Cryptic/Cfc1* (Figures 3B, 5).

Based on the data obtained in *Xenopus*, the three *Tdgf1* paralogs underwent specialization, with *Tdgf1B* (*Xenopus Tdgf1.2*) becoming the exclusive member involved in left-right patterning^7^. *Cryptic/Cfc1* is likewise involved in left-right patterning, suggesting that it has retained its specialization subsequent to the translocation event. This orthologous relationship and conservation of specialization should be further investigated by including analyses from other tetrapod models, in particular reptiles. Reptiles have historically been understudied in developmental biology, but our recent work with veiled chameleons holds a lot of promise for the study of early developmental processes, and roles of orthologous genes in different patterning events^18,19^.

An important area for this field to focus on in the future, is to analyze the consequences of gene loss and its effects on Nodal signaling, like the complete loss of *Tdgf1* in *Ciona intestinalis*. The function of *EGF-CFC* genes in monotremes and marsupials is equally interesting. Both have a single copy of the gene, with platypus retaining the *Tdgf1C* ortholog, and marsupials retaining *Tdgf1B* in its original genomic location. Hypothetically, in both cases the genes could have re-acquired their generalized functions in mesoderm and endoderm formation, as well as anterior-posterior and left-right patterning^2,3^. In contrast, we were not able to identify a *Cryptic/Cfc1* gene in the cow genome and thus whether cow *Cripto/Tdgf1* re-acquired its left-right patterning functions, remains to be determined.

Overall, CFC-EGF proteins have been historically reported to have little sequence conservation between species, beyond the CFC-EGF domain, which remains relatively well conserved. The reasons behind this evolutionary flexibility currently remain unknown.

We observed frequent rearrangements and recombinations in regions proximal to *EGF-CFC* genes, in addition to several duplication and translocation events. The human genome is a good example of the outcomes of these processes, since it contains tandemly duplicated *CFC1* and *CFC1B* genes in the *CRYPTIC/CFC1* locus, and several pseudogenes, including the CRIPTO3 pseudogene on the X chromosome, which resembles CRIPTO (Figure 1B), and contains part of the CRIPTO promoter region^20^. The eutherian locus, containing *Cripto/Tdgf1* has little resemblance to the ancestral *Tdgf1* region due to numerous recombination events, coupled with gene loss. Future analysis should determine whether regions containing *EGF-CFC* genes represent conserved recombination hotspots across different species, and whether proximal transposable elements may explain gene translocations.

In this study we have outlined the evolutionary history of the EGF-CFC family of proteins. We have traced their evolution in deuterostomes from a single gene through their expansion in tetrapods, then specialization, gene loss and translocation in eutherian mammals. EGF-CFC proteins are critical co-factors for Nodal and are thus involved in many processes in early development and pattern formation, and the co-evolution of EGF-CFC and Nodal proteins is remarkable.

## METHODS

### Species phylogenetic tree

We used phyloT (v2 Database information: phyloT database version: 2023.2; NCBI taxonomy nodes: 3 905 559; NCBI GenBank ACCs: 1 007 637 254; RefSeq protein IDs 133 123 611; Uniprot protein IDs/ACCS 332 457 571; Genome Taxonomy Database release 214) (https://phylot.biobyte.de/) to generate the species phylogenetic tree, based on the NCBI taxonomy. Individual species of interest were searched in the NCBI taxonomy to add to the “NCBI tree elements” section. The following tree and file options were selected: internal nodes – expanded; polytomy – yes; file format – Newick; ignore errors – yes. We selected to visualize the tree in iTOL (Interactive tree of life v 6.9)^21^. The tree nodes were rotated to position mammals at the top of the page. Individual scientific names were replaced with common names. For visualization purposes the labels were aligned right, and the scaling factor was set to 0.1 horizontal.

### Gene and protein identification

EGF-CFC genes were first identified using the NCBI Orthologs function^22^. The identity of the genes of interest was ascertained using the neighboring genes in the syntenic block, in comparison to other species. Additionally, we identified some EGF-CFC proteins using BLASTP program, targeting specific species of interest^23^. Lamprey genes and homologous regions were examined using https://simrbase.stowers.org/^14^. Lastly, some of the proteins used in this study had been previously identified by Ravisankar et al.^8^ and Truchado-Garcia et al.^2^

### Protein phylogenetic tree

Individual sequences for Tdgf1/Cripto and Cryptic/Cfc1 homologs were used to determine the relationship of protein sequences. To generate the Tdgf1/Cfc1 phylogenetic tree, we used the https://ngphylogeny.fr/ web interface, selecting “A la Carte” to generate a custom workflow^24^. We selected MAFT for multiple alignment, with “linsi” option selected^25^. We selected trimAI for alignment curation, with 0 gap threshold^26^. For tree inference we selected FastTree with 1,000 bootstrap replicates^27–29^. “Booster Tree with [id avg transfer distances depth] as branch labels” was selected as most representative of both protein relationships and phylogenetic relationships of species of origin. The phylogenetic tree was visualized using iTOL (Interactive tree of life v 6.9)^21^. The scaling factor was set to 0.2 horizontal.

To generate the Fam240 phylogenetic tree, we inputted select protein sequences into Clustal Omega Multiple Sequence Alignment (MSA) program^30^. Phylogenetic tree was exported into iTOL (Interactive tree of life v 6.9) for visualization. The scaling factor was set to 0.2 horizontal.

## Figure editing

Figures were constructed and edited using Adobe Illustrator.

## ACKNOWLEDGEMENTS

We would like to thank members of the Trainor lab for their thoughtful comments and discussion. This work was supported by the Stowers Institute for Medical Research (P.A.T) and a K99 (HD114881) from the National Institute for Child Health and Human Development (N.A.S).

## FIGURE LEGENDS

**Supplementary Figure S1.**
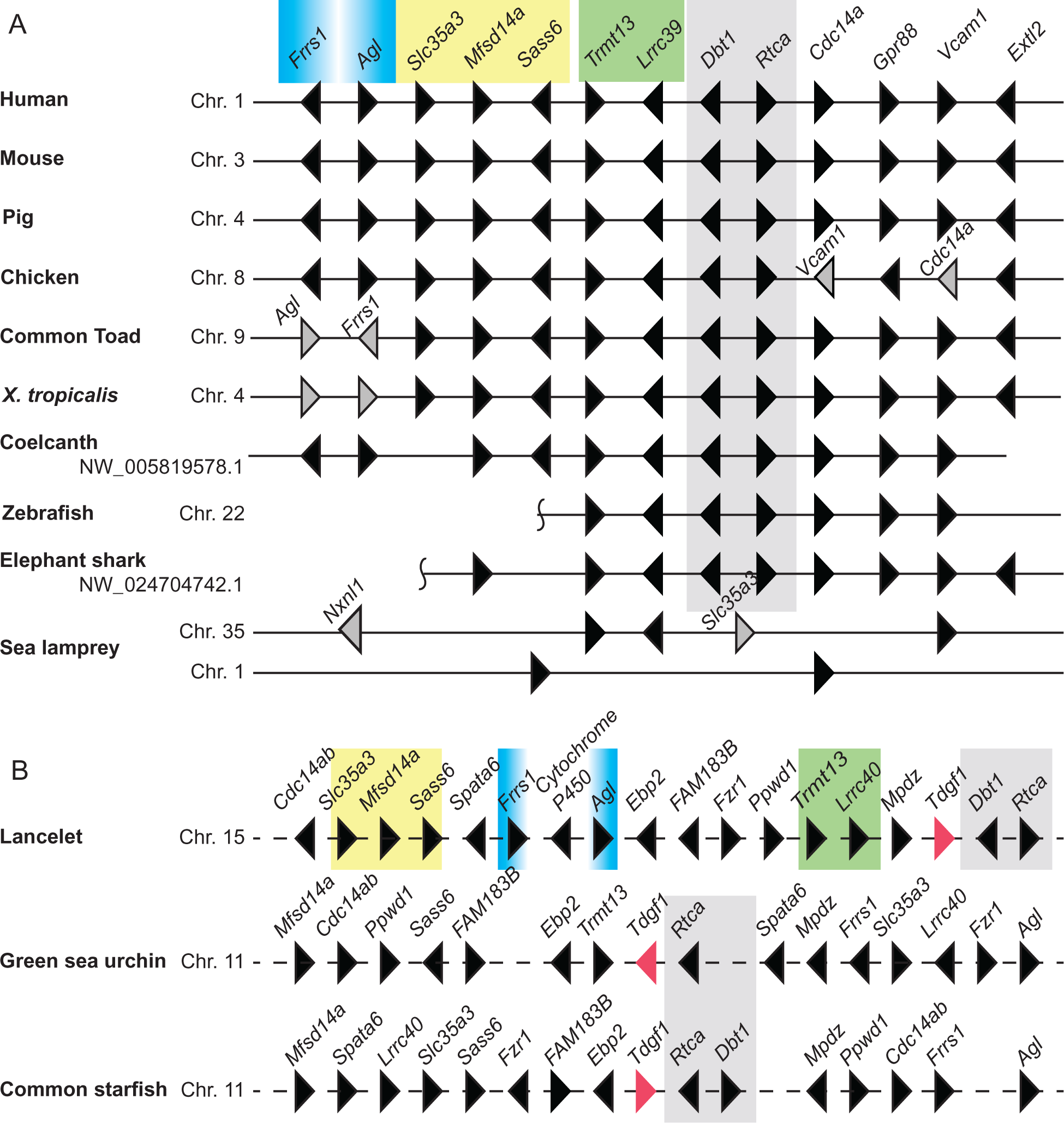
| Genomic regions of homology. A. Structure of the chromosomal region, homologous to *Cripto/Tdgf1* region in craniates. Pictured are key model organisms, and additional species, which aid in revealing conserved regions of homology. **B.** Expanded chromosomal regions and selected landmark genes for lancelet, green sea urchin and common starfish. Only conserved landmark genes are pictured. Additional species-specific genes are present throughout, and not depicted. Ancestral *Tdgf1* gene is colored pink. *Dbt1* and *Rtca* are linked to *Tdgf1* on the ancestral deuterostome chromo-some and are highlighted in gray. Other blocks of linked genes are highlighted in blue, yellow and green. Vertical tildes designate the location of an ancestral chromosomal recombination events. Dashed lines indicate that the genes are on separate distant parts of the same chromosome. Genes with unique minor rearrangements are highlighted in gray and labeled directly on the figure. Chromosome and scaffold numbers are as indicated.

**Supplementary Figure S2.**
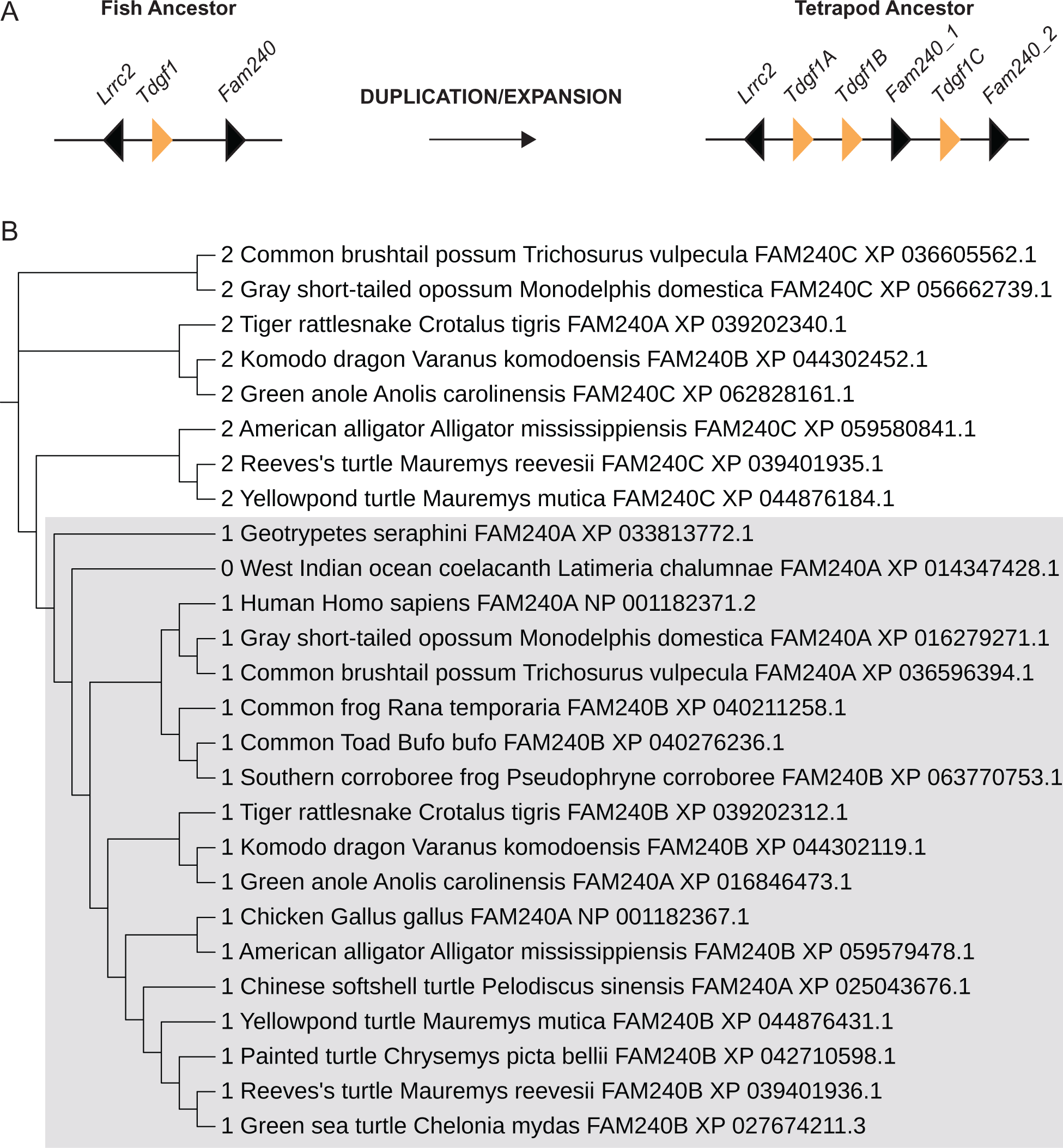
| Expansion of the genomic region containing Tdgf1 and Fam240. A. Diagram showing the expansion of the chromosomal region from a fish ancestor, containing single copies of *Tdgf1* and *Fam240*, proximal to *Lrrc2*, to a tetrapod ancestor, containing three copies of *Tdgf1* (named here A-C, counting from *Lrrc2* gene) and two copies of *Fam240* (named here 1 and 2, counting from the *Lrrc2* gene). *Tdgf1* genes are highlighted in orange. **B.** Phylogenetic relationship of Fam240 protein sequences. For ease of visualization the proteins were labeled with 0 (ancestral), 1 or 2, based on their proximity to the *Lrrc2* gene and relative positioning in respect to *Tdgf1* genes. The orthology group of some of the proteins was determined based on their homology to other proteins. The members of the Fam240_1 orthologous group are highlighted in gray.

